# Transient Early Postnatal Neuronal Hyperexcitation Results in Lasting Social Preference Deficits

**DOI:** 10.1101/2025.09.04.674307

**Authors:** Haleigh Bach, Mikael Nakamura-Vernet, Kevin Lançon, Pierre Drapeau, Hidaya Ben Mhenni, Cameron Oram, Sarah Martin, Philippe Séguéla, Jean-Francois Poulin

## Abstract

Perturbations during critical periods of neurodevelopment are implicated in the etiology of autism spectrum disorder (ASD), a condition marked by considerable heterogeneity and prevalent comorbidities. One such comorbidity is epilepsy, with approximately 30% of children with ASD experiencing seizures and a similar proportion of children with epilepsy displaying ASD-like symptoms. Both conditions have been associated with disruptions in the brain’s excitation-to-inhibition (E/I) balance. Although early-life seizures in rodents have been linked to social impairments, direct causal evidence connecting E/I imbalance, interneuron development, and social behavior remains limited. To address this gap, we induced transient, brain-wide hyperexcitation in neonatal mice using pentylenetetrazol (PTZ), a GABA_A receptor antagonist. We administered both convulsive and subconvulsive doses and assessed long-term effects on social behavior, cortical E/I balance, and parvalbumin (PV) interneuron development. PTZ-treated groups displayed impaired social preference as measured in the 3-chamber test and increased PV interneuron density within the medial prefrontal cortex. These findings highlight a critical developmental window during which E/I imbalance leads to social deficits characteristic of ASD and epilepsy. They also reveal dose-dependent neurobiological changes, underscoring the importance of early-life neural activity in shaping social circuitry.

## Introduction

Brain development progresses through a series of sensitive periods of enhanced plasticity, ranging from embryonic neurogenesis to synaptic pruning in adolescence (Marin 2016; Silbereis et al., 2016). Disruptions during these periods, whether genetic or environmental, are strongly implicated in the etiology of Autism Spectrum Disorder (ASD) (Meredith, 2015). Environmental insults such as pollution, infection, or prenatal stress during the third trimester—especially when combined with genetic susceptibility—have been associated with increased risk of ASD (Mkhitaryan et al., 2025; Love et al., 2024; Raz et al., 2015). Despite its complex etiology, core clinical features such as impaired social behaviour (De Felice et al., 2015; Marin 2016) highlight that diverse insults could converge on common circuit alterations during development. Although the mechanistic underpinnings of ASD remain elusive, its high comorbidity with conditions such as epilepsy could be informative in their shared etiology (Bozzi et al., 2017; Giovagnoli and Smith, 2019). Indeed, epilepsy is more common in children, adolescents, and adults with autism compared to the general population and, depending on the study, co-occurs at a prevalence of up to 30% (Tuchman and Cuccaro, 2011; Liu et al., 2011). Thus, determining if developmental hyperexcitation, or transient imbalances in the E/I ratio, is sufficient to alter social behaviour may explain why ASD is more prevalent in children with epilepsy.

Few studies have assessed whether increasing neuronal activity during a brief developmental window could result in social behaviour impairments. Those that have, however, demonstrated that cortical neuron hyperexcitation during development impairs social behaviour later in life (Bitzenhofer et al., 2021; Medendorp et al., 2021). For instance, one study used Emx1-Cre mice and a bioluminescence-coupled chemogenetic approach to activate cortical pyramidal neurons from postnatal days 4-14 (P4-14), a broad window corresponding to both the pre- and postnatal period in humans (Medendorp et al., 2021; Semple et al., 2013). In adulthood, these mice exhibited reduced social preference in the three-chamber test. In contrast, Bitzenhofer and colleagues (2021) used optogenetics to stimulate layer 2/3 (L2/3) neurons of the cortex from P7-11, resulting in impaired social preference at P21, which persisted into early adulthood. Notably, this window overlaps with the maturation of parvalbumin-expressing interneurons (PVINs), which follows a wave of cortical interneuron apoptosis occuring between P5 and P10 in mice (Lim et al., 2018; Wong et al., 2018). PVINs are essential to proper circuit formation and function as their fast-spiking properties generate gamma oscillations necessary for the development and refinement of cortical networks (Cardin et al., 2019; Denaxa et al., 2018; Miyamae et al., 2017; Sohal et al., 2009).

PVINs are also known to be altered in function and amount in ASD and epilepsy (Jiang et al., 2016; Contractor et al., 2021). Postmortem studies have revealed reduced PVIN density in the medial prefrontal cortex (mPFC) of individuals with ASD (Hashemi et al., 2016; Varghese et al., 2017), and mouse models such as the *Shank1*^-/-^ and *Shank3b*^-/-^ mutants (Felice et al., 2016). In addition, several epilepsy-associated genetic mutations, including those affecting the transcription factor *Arx*, disrupt PVIN development (Jiang et al., 2016; Marsh et al., 2009). Further work also highlights the role of PVINs in regulating social behaviour with one study identifying a specific mPFC PVIN subpopulation that was sufficient for initiating and maintaining social approach behaviour in mice (Bicks et al., 2020). Optogenetic activation of these interneurons enhanced social interactions, while chemogenetic inhibition reduced time spent in social zones during the three-chamber test. Thus, PVINs are not only altered in ASD and epilepsy, but are crucial for generating appropriate social responses in adulthood.

While previous literature provides insight into the role of developmental hyperexcitation in social behaviour impairments, there are important gaps that need addressing. For example, it is unclear whether convulsive hyperexcitation elicits a more severe phenotype than non-convulsive hyperexcitation (Bitzenhofer et al., 2021; Castelhano et al., 2013; Medendorp et al., 2021; Lugo et al., 2014). This is an important distinction worth investigating, given individuals with ASD experienced increased infantile spasms and early-life seizures compared to the general population without necessarily meeting diagnostic criteria for epilepsy (Tuchman and Cuccaro, 2011). To address these gaps, we administered the GABA_A_ receptor antagonist pentylenetetrazol (PTZ) in mice from P8-11 and P11-14, to determine if there was a window in which social behaviour was most susceptible to hyperexcitation. Additionally, we assessed if social behaviour, PVIN formation, and intrinsic cortical activity varied by whether PTZ was administered at a convulsive or non-convulsive dose. We found P8-11 to be a window in which social behaviour was impaired, but the social behaviour impairment did not vary by dosage. However, PVINs formation and intrinsic mPFC pyramidal neuron activity had dosage-specific variations. The findings of the present study shed light on the developmental mechanisms underlying behavioural and neuronal phenotypes relevant to the co-occurrence of ASD and epilepsy.

## Methods

### Mouse Colony Management and Neonatal Care

All procedures involving the use of animals were approved by the Montreal Neurological Institute Animal Committee (AUP 8132). Mice were housed with a 12-hour light-dark cycle with a constant temperature and humidity, and *ad libitum* access to food and water. All procedures were conducted during the light phase of the light-dark cycle between 7:00am and 7:00pm. C57BL/6 (wildtype) mice were the only type of mouse used in this study and were from the Jackson Laboratory. Experiments conducted on pups aged P1-14 involved separation from the dam into a transfer cage. When separated, the entire litter was removed at the same time and rubbed with bedding prior to being returned to the home cage to control for any effects of maternal stress and decrease the chance of maternal rejection (Chen et al., 2024).

### PTZ Injections

Mice were intraperitoneally injected with 30 or 50mg/kg of pentylenetetrazol (PTZ) from P8-11 or P11-14. The 30mg/kg dose has been used in previous studies of chemokindling in which several doses were administered per day over several days to induce epilepsy (Monteiro et al., 2024; Shimada and Yamagata, 2018). To our knowledge, the 50mg/kg dose is the lowest dose of PTZ reported to induce generalized seizures in neonates with a single injection per day over four days (Monteiro et al., 2024; Parker et al., 2016). Injections were conducted according to the protocol of Porcratsky and Sleigh (2023) with the adaptation of using a 0.03ml, 31G insulin syringe. Neonates were weighed on each injection day to calculate the appropriate volume of PTZ or saline for the dose.

### Recording Evoked and Spontaneous Neuronal Activity in Slices

Both the evoked and spontaneous post-synaptic currents of anterior cingulate cortex (ACC) layer 2/3 pyramidal neurons were measured with whole-cell voltage clamp electrophysiology in acute mouse brain slices. Recordings were taken from P32-40 PTZ-injected mice following behavioural testing. Mice were anesthetized with an Avertin solution (2.5 g tribromoethanol in 5 mL amylene hydrate diluted in 100 mL ddH2O) (Sigma-Aldrich, St. Louis, MI), then transcardially perfused with a 4°C choline-chloride based cutting solution oxygenated with carbogen (O_2_ 95%, CO_2_ 5%). Brains were extracted and sliced into coronal sections of 300 µm using a vibratome (Leica VT1000). Slices rested at room temperature for 1 hour in oxygenated artificial cerebrospinal fluid (aCSF) containing 124 mM NaCl, 2 mM KCl, 26 mM NaHCO3, 1.8 mM MgSO4, 1.25 mM NaH2PO4, 10 mM Glucose, 1.6 mM CaCl2, pH 7.4. Slices were submerged at 30-32°C on the stage of a Zeiss Axioskop microscope continuously perfused with oxygenated aCSF. Cortical pyramidal cells were identified by morphology and were patched with glass pipettes (∼ 6 MΩ) filled with an intracellular solution containing 125 mM CH3O3SCs, 10 mM NaCl, 2 mM MgCl2, 10 mM HEPES, 2 mM Mg-ATP, and 0.4 mM Na-GTP. IPSC and EPSC recordings were taken from L2/3 cells by holding them at -70mV or -10mV, respectively. Synaptic events were analyzed using MiniAnalysis (Synaptosoft).

### Transcardiac Perfusions

Mice between P30-80 were deeply anesthetized in an isoflurane chamber with an O_2_ flow rate of 2 L/min and 5% isoflurane. Mice were then transcardially perfused with phosphate-buffered saline (PBS) then cold 4% paraformaldehyde (PFA) in PBS. If mice were perfused between P11-14, the procedure was modified with a reduced volume of solution and a slower flow rate of perfusion (Perez Arevalo et al., 2022). Brains were then dissected, fixed overnight in 4% PFA, then cryoprotected in 30% sucrose in PBS for 48 hours. Brains were then frozen in optimal cutting temperature compound (OCT) at –80°C until they were sectioned at 25μm using a cryostat.

### Immunostaining

Immunofluorescence was conducted through rinsing sections with PBS, blocking in 5% normal donkey serum in PBS + Triton-X-100 (0.3%), and incubating with primary antibodies to detect PVINs (1:2000, Millipore Sigma, catalogue number MAB1572), Wisteria floribunda agglutinin (WFA) to mark PNNs (1:500, Millipore Sigma, catalogue number L1516), or cFos (1:100, Cell Signalling Technology, catalogue number 31254S). The antibodies were diluted in blocking buffer and brain sections incubated overnight at 4°C. The following day, the sections were rinsed in PBS + Tween-20 (0.05%) and incubating with secondary antibodies (anti-mouse AlexaFluor 488, Thermo Fisher, catalogue number A-21202; anti-rabbit AlexaFluor 488, Thermo Fisher, catalogue number A-21206; streptavidin, AlexaFluor 647 Conjugate, Thermo Fisher, catalogue number S21374) diluted in blocking buffer with DAPI (1:1000) to mark nuclei. Sections were rinsed with PBS + Tween-20 0.05% and then PBS prior to being mounted onto slides and left to dry. Once dry, the slides were coverslipped with Gelvatol and stored at 4°C prior to imaging.

To assess cellular activity, the immediate early gene marker, cFos, was used. Mice were injected with 30 or 50mg/kg of PTZ from P8-11 and perfused approximately 1-1.5 hours post-injection on P11 to allow for maximal protein expression (Kovács 1998). Brains were processed as described above, with the addition of antigen retrieval to facilitate binding to the epitope. This involves immersing the slides in a sodium citrate buffer (10mM Sodium Citrate, 0.05% Tween 20, pH 6.0) that is pre-heated to 80°C Celsius in a water bath for 30 minutes (Jiao et al., 1999). The slides were then removed from the solution, left to cool for 10 minutes, then rinsed with PBS + Tween 20 (0.05%). and put into blocking buffer to continue the staining as described above.

### Microscopy and Cell Quantification

Images were acquired using the Nikon Ti2 confocal microscope at 10x (PVINs) and 20x (cFos) magnification. Confocal images of the mPFC (combined IL and PL), ACC, Caudoputamen (CPu), somatosensory cortex (S2), and hippocampus (CA1) were captured using the same laser parameters between treatment and controls. Cell counts were obtained using NIS ai on the NIS Elements analysis software. A unique NIS ai was trained to count cells positive for cFos, parvalbumin, and WFA. Training images included both treatments and controls in each brain region and training was conducted until the error rate was below 0.05%.

For PVIN quantifications, L2/3 and L5/6 of mPFC and ACC images were identified using the Bigwarp plugin on FIJI that we used to match landmarks from fixed (confocal) images to brain sections from the Allen Reference Atlas (Bogovic et al., 2016). The warped atlas images were used to trace of the cortical regions and layers on NIS-AR. The CPu, S2, and CA1 were detected using NS ai trained to recognize these regions. All automatically determined ROIs were screened prior to performing cell counts and manually corrected if found inaccurate. Cell counts for each ROI were conducted with the trained NIS ai.

### Behavioural Tests

#### Three-Chamber Test

The three-chamber test was used to measure social preference and social novelty preference in 4–5-week-old mice following a modified protocol published by Rein et al., (2020). All behavioural tests were conducted in dark conditions with the experimenter in a separate room to limit stress. Our tests included habituation to the chamber (phase one), a social preference test (phase two), and a social novelty test (phase three). We did not include the pre-test component with the paper balls placed under the cups. Prior to all tests, mice were habituated to the behaviour room for 30 minutes in their home cage in the dark. Test mice were also habituated to the three-chamber apparatus without stimuli for two days before the testing day. The stranger mice were habituated to the cups for two days without test mice.

Mice in the three-chamber recordings were tracked using DeepLabCut (Mathis et al., 2018). A custom Matlab script was used quantify frames that showed close interactions between experimental mice and the stimuli. DeepLabCut tracking data could then be used in custom MatLab script to generate outputs for percent interaction time and images of the animal’s track overlayed on video frames. The percentage of time interacting with either stimulus in phase two and three determined social preference and novelty preference, respectively. Social preference indices were calculated as the time of interaction with social stimulus over the total interaction time.

#### Open Field and Free Dyadic Social Interaction Tests

We combined the open field and free dyadic social interaction tests into one test spanning 10 minutes. This test was conducted in a modified square-shaped open field arena measuring 40x40cm. These tests were run on mice between P27-33 either the day prior to, or following, the three-chamber test. The open field test was conducted during the first five minutes in which the mouse could freely explore the arena and followed the protocol of (Seibenhener and Wooten 2015). Following those initial five minutes, a sex-matched, age-matched stranger mouse was added to the center of the arena for the free dyadic interaction test. The test mouse and novel conspecific were left for five minutes and we followed the protocol described by Kraeuter et al., (2019).

Video recordings from the open field and free dyadic interaction tests were manually annotated by a researcher blinded to the conditions with a custom ethogram (Table 1) using the Behavioural Observation Research Interaction Software (BORIS) software (Friard and Gamba, 2016). For the first five minutes (i.e., in the open field test), instances of self-grooming were manually scored. Animals were tracked using Social LEAP estimates animal poses software (SLEAP, Pereira et al., 2022). Custom Python code was used to pre-process the SLEAP tracking data by removing unwanted instances, adding non-existing frames, and accounting for track switches between the animals. Afterwards, the x, y coordinates were transformed to fit a common 40 by 40 cm field to account for different recording angles, heights and positions. The center of the arena was defined as the middle 40% of the area, as done in previous (Peixoto et al., 2019; Seibenhener and Wooten 2015). Data from SLEAP were also used in a custom MatLab code to generate normalized heatmaps for each group on the open field arena.

For the free dyadic social interaction test, nose-to-nose interactions and instances of play were manually annotated with BORIS. These videos were run through SLEAP for animal tracking using the same model from the open field test. BORIS was also used to identify/correct track identity switches during dyadic interactions. Output of animal tracking data from SLEAP was used in custom Python code to quantify instances of nose-to-nose interactions. To ensure that the automated quantifications of interactions used in our data analysis were accurate, we compared them to interactions manually quantified from BORIS.

#### Von Frey Test

The von Frey test was used to assess mechanical sensitivity in P33-40 mice injected with 30 or 50mg/kg of PTZ from P8-11 (Deuis et al., 2017). This was conducted with at least a one-day break following the sociability testing. Male and female mice were separately habituated to- and tested on the apparatus. Mice were habituated to the apparatus without the microfilament for two hours on the first day and one hour on the second day. On testing day, mice were placed on the apparatus for 30 minutes for short-term habituation. Then, withdrawal thresholds were determined by perpendicularly poking the lateral planar surface of both hindpaws with an automatic von Frey apparatus (Ugo Basile) and recording the threshold. Three trials were performed for each paw with a break of at least two minutes between stimulations. Six withdrawal threshold values were recorded and averaged for each mouse.

### Statistical Analysis

All data were analysed using GraphPad Prism 10. The values shown are the means ± SEM. Significance threshold was p<0.05. For t-tests, the Rout correction verified there were no outliers. All analyses were also run with sexes separated if the sample size was greater than three. If analyses on sex differences were not included in a figure, this means there was no sex-specific differences Some graphics and diagrams were created with BioRender and figures were assembled with Affinity Designer.

## Results

### PTZ Injections Between P8-11 in Mice Reveal a Critical Window for the Development of Social Behaviour

To examine the effect of early life neuronal excitation on social behaviour, mice were injected with subconvulsant (30mg/kg) doses of PTZ or saline between P8-11 (Fig. 1*A*). Social preference was assessed at P30 by comparing the percentage of time spent interacting with an object versus a conspecific for 10 minutes (phase 2). A repeated-measures two-way ANOVA yielded a significant main effect of PTZ on percentage interaction time (effect of stimulus F(1, 26) = 20.06, p = 0.0001; interaction effect F(1,26) = 3.48, p = 0.0735; Saline n = 13, PTZ n = 15; Fig. 1*C*). While saline-injected mice displayed a significantly increased percentage of time interacting with the mouse over the object (p = 0.0004) whereas this preference was absent in PTZ-injected mice.

**Figure 1.**
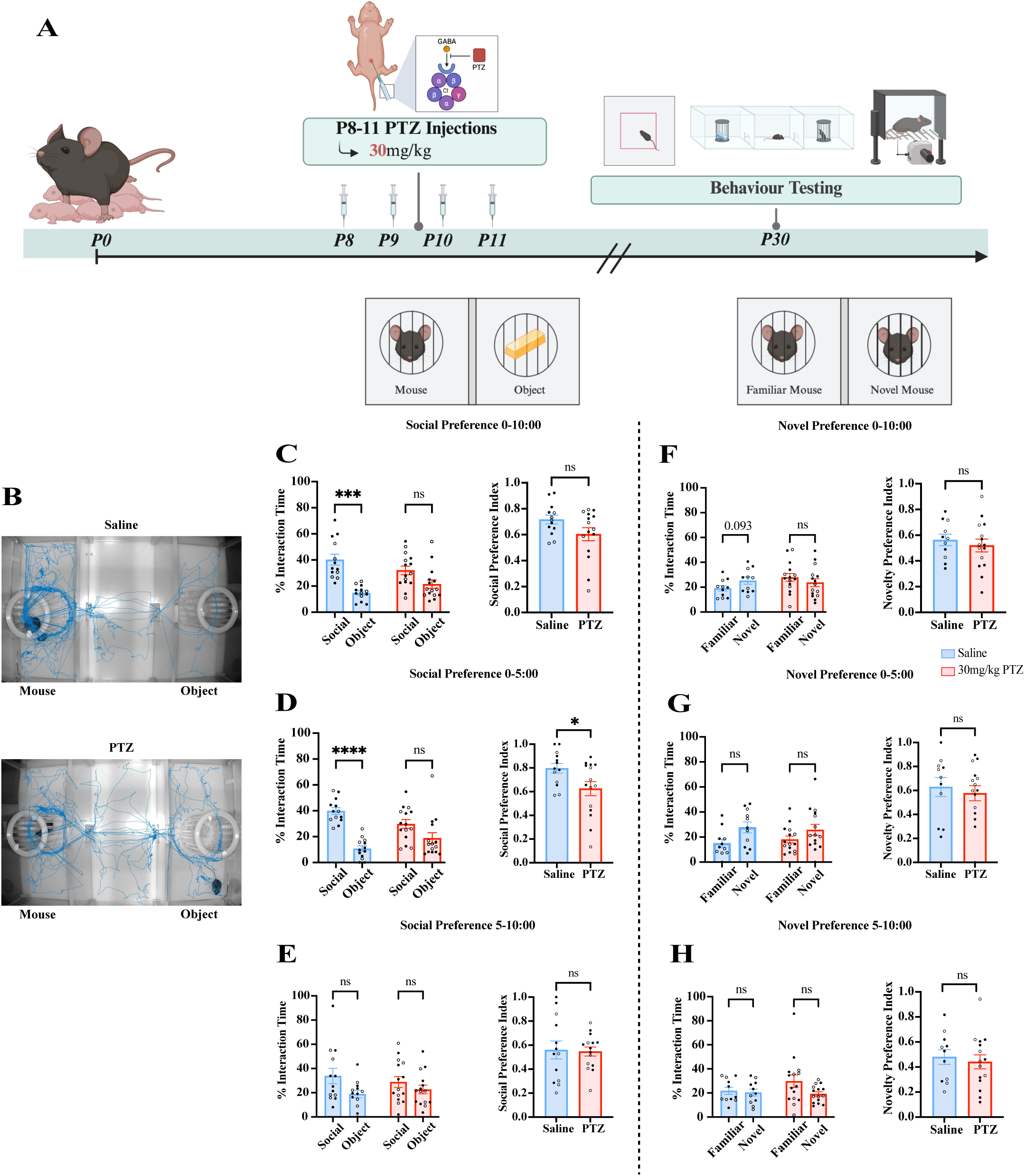
Subconvulsive PTZ treatment at P8-11 alters social preference in the three-chamber test at P30. ***A***, Experimental overview of P8-11 30mg/kg PTZ injections and behaviour testing at P30. ***B***, Representative tracking of saline-(top) and PTZ-injected (bottom) mice during phase two. PTZ-injected mice displayed a lack of significant social preference compared to controls during the full time (***C***), first half (***D***), but not the last half (***E***) of phase two. ***F-H,*** PTZ-injected and control mice do not display a significant preference for novel interaction throughout phase three. Bars represent mean ± SEM. Points represent individual mice. Filled circles represent females while unfilled circles represent males. P-values determined by a two-way ANOVA with the Bonferroni correction or unpaired Student’s t-tests. *p < 0.05, ***p < 0.001, ****p < 0.0001.

Previous studies analyzing sociability in the three-chamber test have found that social preference is strongest during the initial minutes of the phase (Nadler et al., 2004; Page et al., 2009). As these studies observed this effect in adult mice, we were curious whether this pattern would emerge in P30 mice. To explore this, we divided phase two into the first and last five minutes. We found a significant interaction effect between treatment and stimulus for the first half (Interaction effect F(1,26) = 4.94, p = 0.0351; Fig. 1*D*), indicating that the proportion of interaction time varied depending on both treatment (PTZ or saline) and stimulus. Additionally, there was a significant main effect of stimulus (social vs. non-social) on percentage interaction time during the first five minutes (effect of stimulus F(1, 26) = 23.90, p < 0.0001), with only saline-injected mice displaying a social preference (p < 0.0001). The social preference index of the first half corroborated this finding, as saline-injected mice exhibited a significantly increased ratio of social interactions compared to the PTZ-injected mice (Unpaired Student’s t-test t(26) = 2.306, p = 0.0294; Fig. 1*D*). There were no statistically significant differences in social preference index or percentage interaction time for either group in the last five minutes of phase two (Fig. 1*E*). This finding suggests that social preference is strongest during the beginning time intervals of the three-chamber test in juvenile mice treated with saline, aligning with what was shown in adults (Nadler et al., 2004; Page et al., 2009).

We then investigated whether juvenile mice exhibited a preference for novel social interactions, a behaviour typically observed in wildtype mice as young as six weeks old (Moy et al., 2004), and if this preference was impaired by P8-11 PTZ-injections. We did not observe a significant preference for a novel mouse over a familiar one, in neither saline-nor PTZ-injected mice at P30 (Fig. 1*F-H*). There were also no significant differences in novel preference indices between treatments and controls. This lack of significant novelty preference observed in control prepubescent mice contrasts with previous reports in adult mice, which typically exhibit a preference for novel social stimuli (Gunaydin et al., 2014; Moy et al., 2004). This suggests that the preference for novel social stimulus may not yet be established in juvenile mice at P30. Taken together, subconvulsant PTZ injections during P8-11 developmental window affect social preferences in P30 mice.

### The Behavioural Phenotype Induced by P8-11 PTZ Injections is Specific to Social Preference

We next wanted to examine additional behavioural alterations resulting from PTZ injections. We first aimed to characterize social behaviour effects in a more naturalistic setting by conducting a modified free dyadic social interaction test (Fig. 2). The open field and free dyadic social interaction tests were combined into one test where a mouse was first alone in an open field arena for five minutes before a novel mouse was introduced. We quantified instances of grooming (as a measure of repetitive behaviour), distance travelled (locomotor/exploratory effects), and time in the center versus periphery (anxiety-like behaviour). We found no significant differences in total time grooming or average duration of grooming bouts between the PTZ- and saline-injected mice (Fig. 2*B*, *C*). Similarly, no significant differences were found in the proportion of time spent in the center of the arena between groups (Fig. 2*D*, *E*). Additionally, there were no significant differences in distance travelled throughout the arena (Fig. 2*G*). We observed no differences between sexes for any of these measures. These results indicate that subconvulsive PTZ treatment from P8-11 did not increase anxiety-like behaviour, repetitive behaviour, or activity levels.

**Figure 2.**
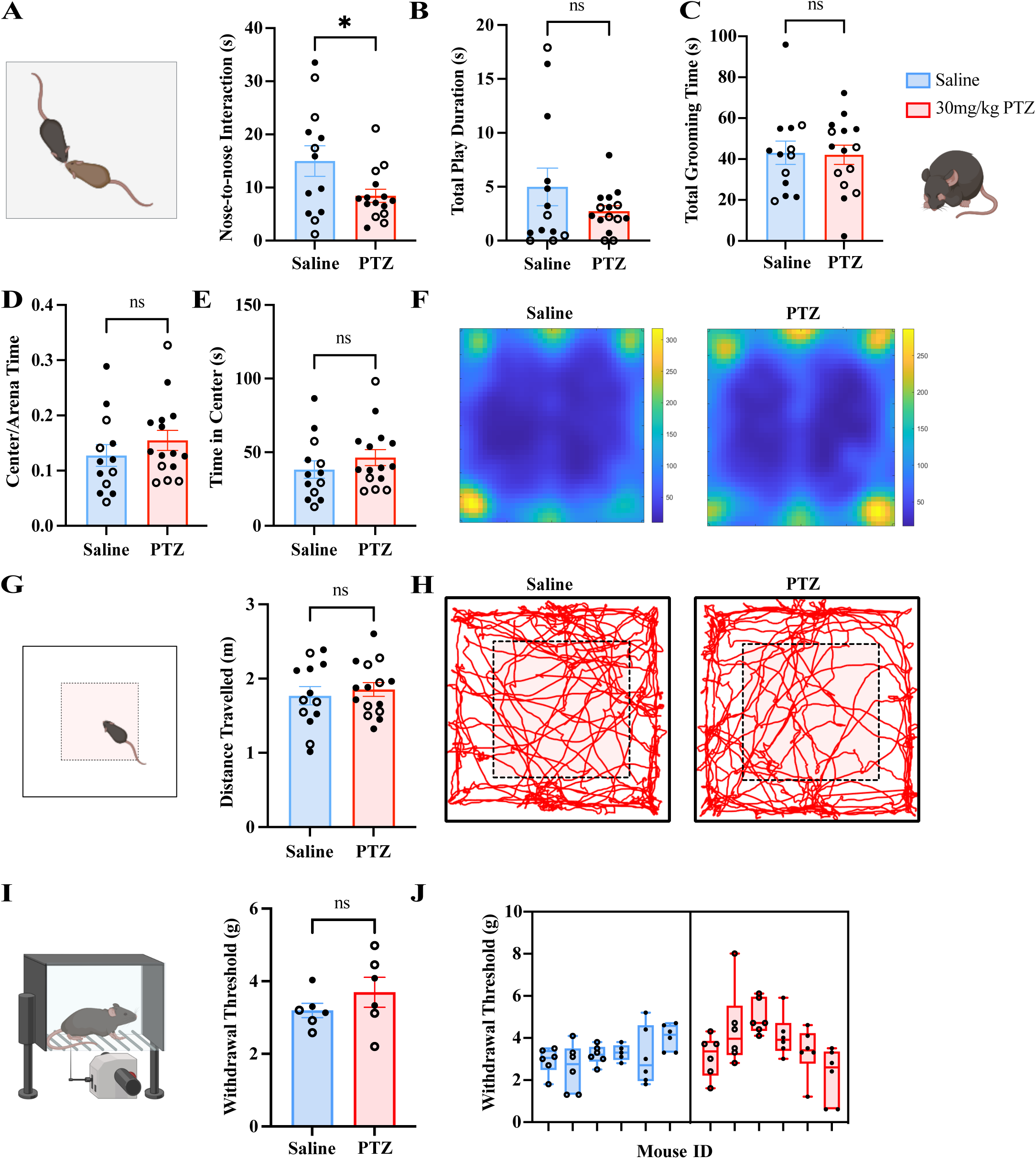
Subconvulsive PTZ treatment at P8-11 reduces interactions without altering anxiety, locomotion, repetitive behaviour, or mechanical sensitivity at P30. ***A***, PTZ-injected mice displayed a significantly reduced nose-to-nose interactions compared to saline-injected mice. ***B***, PTZ-injected mice did not display any significant differences in total play duration. ***C***, PTZ-injected mice did not display any significant differences in total grooming duration. PTZ-injected mice did not display any significant differences in proportion of time spent in the center of the open field arena (***D***) or total time in the center **(*E*). *F***, Representative heat maps of time spent within the open field arena of saline-(left) and PTZ-injected (right) mice. ***G***, There were no significant differences in distance travelled in the open field test. ***H***, Representative tracking of mice the open field arena of saline-(left) and PTZ-injected (right) mice. ***I***, There were no significant differences in withdrawal threshold on the Von Frey test between PTZ- and saline-injected mice. ***J***, Side-by-side boxplots depicting withdrawal thresholds of individual trials in each mouse. Bars represent mean ± SEM. Points represent individual mice. Filled circles represent females while unfilled circles represent males. P-values determined by unpaired Student’s t-tests. *p < 0.05.

After the first five minutes of the open field test, an age- and sex-matched stranger mouse was introduced in the arena, and mice were allowed to interact for five minutes. We then quantified juvenile play and nose-to-nose interactions (Fig. 2*A*, *B*). We found no significant differences average duration of play behaviour between groups (Fig. 2*B*) but observed a decreased number of nose-to-nose interactions in PTZ-injected mice as compared to the saline-injected mice (Unpaired Student’s t-test t(26) = 2.196, p = 0.0372; Saline n = 9, PTZ n = 13; Fig. 2*A*). There were no significant differences in play or between males and females.

Sensory abnormalities are frequently reported in ASD preclinical models and patients. To evaluate whether early-life hyperexcitation affects mechanical sensitivity, we conducted the von Frey test between P32-40. The von Frey assay measures responses to tactile stimulations by applying increasing pressure from an automatic filament on the hindpaws until a withdrawal threshold is reached. Withdrawal thresholds did not significantly differ between PTZ-injected mice and controls (Fig. 2*I*). However, there was considerable variability between and within groups of mice for each of the six trials (Fig. 2*J*). Thus, our data indicate that the effects of PTZ-induced neuronal hyperexcitation from the sensitive window of P8-11 alters the development of social preference and social interaction without altering other behaviours like anxiety, grooming, locomotion, or mechanical sensitivity.

### The Sensitive Window for the Development of Social Preference May Extend beyond P11

Having established P8-11 as a critical period for the development of social preference, we next aimed to determine if social preference could be altered in a different developmental window. Thus, we performed PTZ injections in mice from P11-14, then conducted the three-chamber test at P30 (Fig 3*A*). A two-way repeated-measures ANOVA revealed a significant main effect of stimulus on percentage interaction time for the P11-14 cohort (effect of stimulus F(1,12) = 10.39, p = 0.0073; interaction effect F(1,12) = 0.03, p = 0.8713, Saline n = 8; PTZ n = 6; Fig. 3). Similar to our P8-11 findings, the P11-14 saline-injected mice displayed a significant preference for interacting with the conspecific over the object (p = 0.0475). By contrast, PTZ-injected mice did not display a social preference. However, when the phase was divided into the first and last five minutes, there were no significant differences in the percentage of time interacting with the mouse or the object for the PTZ- or saline-injected mice. This contrasts with the findings of the P8-11 saline group, which displayed a significant social interaction preference in the first half. In phase three, we did not observe a significant preference for novel, or familiar, social interactions in saline- or PTZ-injected mice for the entire testing period (Fig. 3). Additionally, when phase three was analysed when divided by the first and second halves, there were no significant preferences for novel or familiar interactions detected in either the saline- or PTZ-injected mice of either cohort. The lack of social novelty preference in this additional group of saline-injected mice suggests a social novelty preference may not be detectable at P30. In summary, these data show that daily subconvulsant PTZ injections between P11-14 induce a sociability deficit, suggesting the vulnerability window may extend beyond P11.

**Figure 3.**
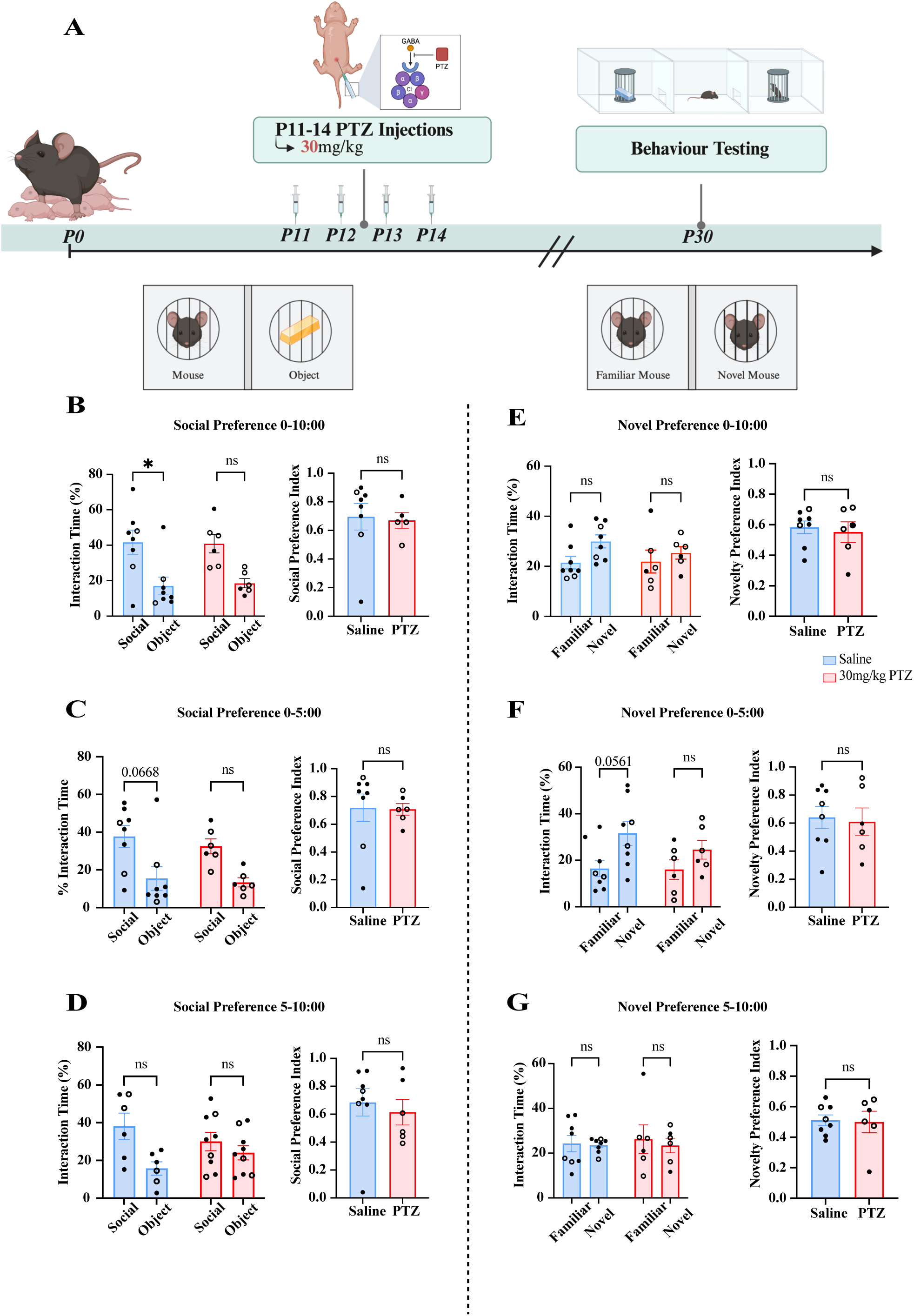
Subconvulsive PTZ treatment at P11-14 alters social preference in the three-chamber test at P30. ***A***, Experimental overview of P11-14 30mg/kg PTZ injections and the three-chamber test at P30. PTZ-injected mice displayed a lack of significant social preference compared to controls during the full time (***B***), but not first (***C***) or second half (***D***) of phase two. ***E-G,*** PTZ-injected and control mice do not display a significant preference for novel interaction throughout phase three. Bars represent mean ± SEM. Points represent individual mice. Filled circles represent females while unfilled circles represent males. P-values determined by a two-way ANOVA with the Bonferroni correction or unpaired Student’s t-tests. *p < 0.05.

### Transient Subconvulsive and Convulsive PTZ Doses Have Similar Effects on Neuronal Activity and Behaviour

Given that 30mg/kg of PTZ is a subconvulsive dose, we wondered if a higher dose would exacerbate the impact on behaviour and assessed the effect of the minimally convulsive dose of 50mg/kg (Monteiro et al., 2024; Parker et al., 2016; Shimada and Yamagata, 2018). We first determined whether 30- and 50mg/kg PTZ injections were sufficient to alter neuronal activity by performing immunofluorescence staining of the immediate early gene cFos (Fig. 4*A*, *B*). The staining was conducted on brains from P11 mice, 1-1.5 hours post-injection to allow cFos protein expression (Kovács 1998), following the same P8-11injection regimen described previously. We quantified cFos-positive nuclei in five brain regions selected for their involvement in social behaviour (Fig. 4*B*): the mPFC and ACC, which are implicated in social behaviour; the mPFC, CPu, and S2, which exhibit altered E/I balance in models of ASD; and the hippocampal CA1 region, which exhibits altered development following early life seizures (Antoine et al., 2019; Bozzi et al., 2017; Greenhill and Jones 2010; Lee et al., 2016; Steif et al., 2007). Quantification of confocal images revealed that mice treated with 30mg/kg PTZ displayed an increased number of cFos positive nuclei in all brain regions analysed compared to controls (effect of treatment F(1,4) = 23.67, p = 0.0083; effect of brain region F(4,16) = 14.51, p < 0.0001; interaction effect F(4,16) = 2.16, p = 0.1199; Saline n = 3, PTZ n = 3; Fig. 4*C*). Bonferroni-corrected post-hoc comparisons showed significantly increased cFos-positive nuclei in the PTZ group in the mPFC (p = 0.0171), ACC (p = 0.0007), CPu (p = 0.0135) and S2 (p < 0.0001). The increase in CA1 was trending towards, but did not reach statistical significance (p = 0.0797). The 50mg/kg group showed a similar increased in the number of cFos positive nuclei in cortical brain regions (Two-way repeated-measures ANOVA, interaction effect F(4,16) = 3.14, p = 0.0437; effect of treatment F(1,4) = 18.21, p = 0.0130; effect of brain region F(4,16) = 11.38, p = 0.0001; Saline n = 3, PTZ30 n = 3; Fig. 4*D*). Post-hoc Bonferroni tests showed significant increases in cFos-labelled nuclei in the mPFC (p = 0.0008), ACC (p = 0.0005), and S2 (p = 0.0052) in PTZ-injected mice compared to controls. Overall, these data confirmed that both subconvulsive and convulsive PTZ treatment from P8-11 lead to increased cortical neuronal activity at P11 compared to saline controls.

**Figure 4.**
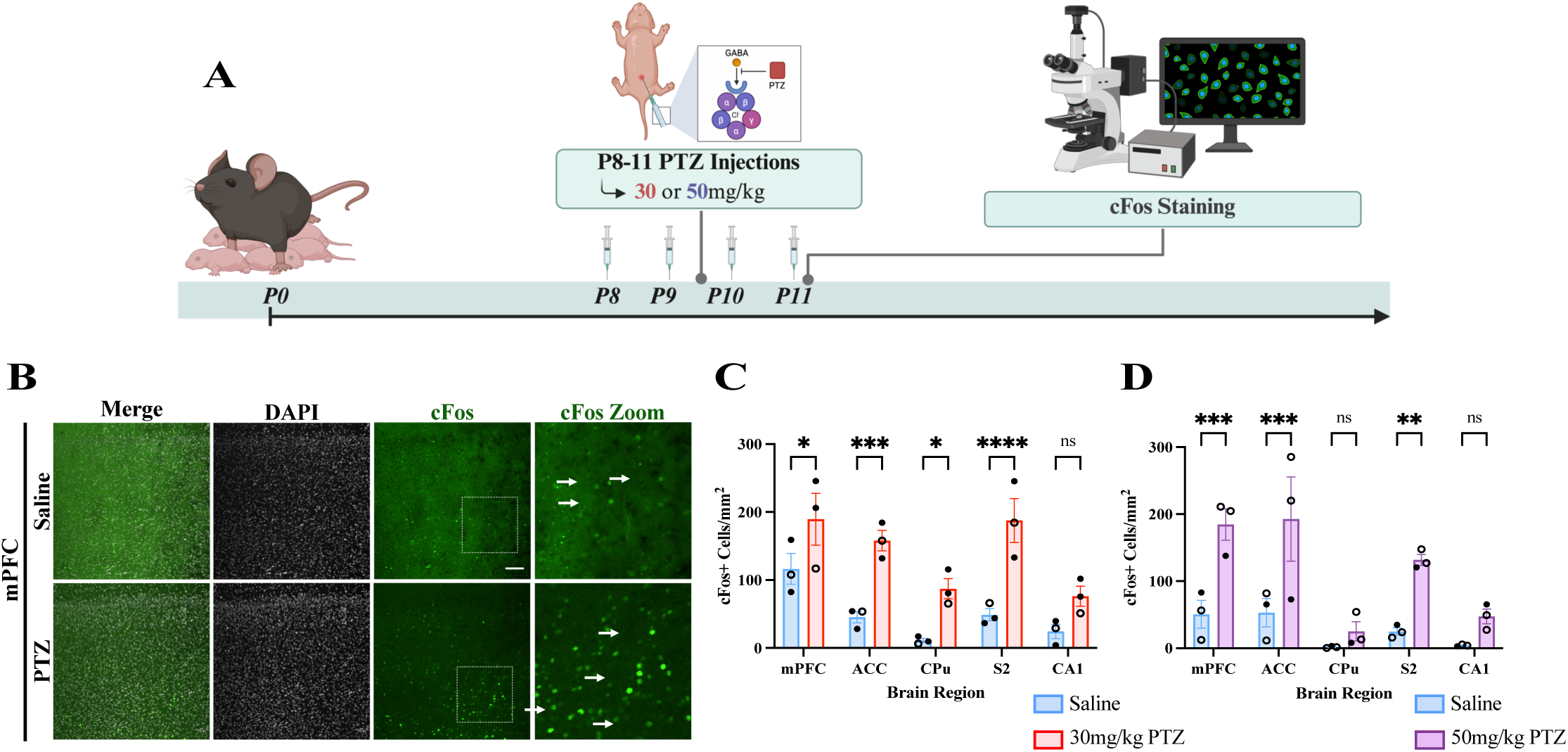
Both subconvulsive and convulsive PTZ treatments increase cortical cFos+ cell density at P11. ***A***, Experimental overview of P8-11 PTZ injections and cFos staining at P11. ***B,*** Representative images of cFos staining in 30mg/kg PTZ- and saline-injected mice at P11 in the mPFC. ***C,*** 30mg/kg PTZ-injected mice display significant increase in cFos+ cell density in the mPFC, ACC, CPu, and S2. ***D,*** 50mg/kg PTZ-injected mice display significant increase in cFos+ cell density in the mPFC and ACC, and S2. Scale bar is 100 µm. Arrows point towards cFos-positive nuclei. All figures represented by mean ± SEM with individual mice as points. Filled circles represent females while unfilled circles represent males. P-values determined by a two-way ANOVA with the Bonferroni correction. * p < 0.05, **p < 0.01, ***p < 0.001, ****p < 0.0001.

To determine whether PTZ treatment induced long-term changes in neuronal activity, and if this is dose-dependent, we performed whole-cell voltage clamp recordings of pyramidal neurons in layer 2/3 of the ACC (Fig. 5*A*). Measurements of spontaneous activity, including frequency and amplitude of spontaneous EPSCs (sEPSCs) and sIPSCs, were collected. There were no significant differences in amplitude of sEPSCs or sIPSCs between the 30mg/kg, 50mg/kg, and saline groups (Fig. 5*C*, *E*). As a change in EPSC and IPSC amplitude is correlated with post-synaptic expression of AMPA and GABA_A_ receptors, respectively, this indicates that PTZ treatment might not have affected the postsynaptic density of these receptors (Kilman et al, 2002). However, mice injected with 50mg/kg PTZ displayed a significantly reduced sEPSC frequency compared to controls and to the 30mg/kg PTZ group (Two-way ANOVA, effect of treatment F(2,67) = 6.5, p = 0.0026; Saline vs PTZ50: p = 0.0079, PTZ30 vs 50: p = 0.0063; Saline n = 49 cells, PTZ30 n = 36 cells, PTZ50 = 33 cells; Fig. 5*B*). There were no significant differences in sIPSC frequencies between groups (Fig. 5*D*). These data illustrate that early-life hyperexcitation with convulsive PTZ from P8-11 reduces spontaneous synaptic activity in layer 2/3 pyramidal neurons at P37-40. Interestingly, subconvulsive 30mg/kg PTZ treatment may not induce long-term changes in neural activity despite increased cortical cFos activation at P11 (Fig. 4*C*). Overall, these results demonstrate the long-lasting effects of convulsive P8-11 PTZ injections in the ACC, however, it is possible that subconvulsive PTZ injection could have affected other cortical regions involved that modulate social behaviour.

**Figure 5.**
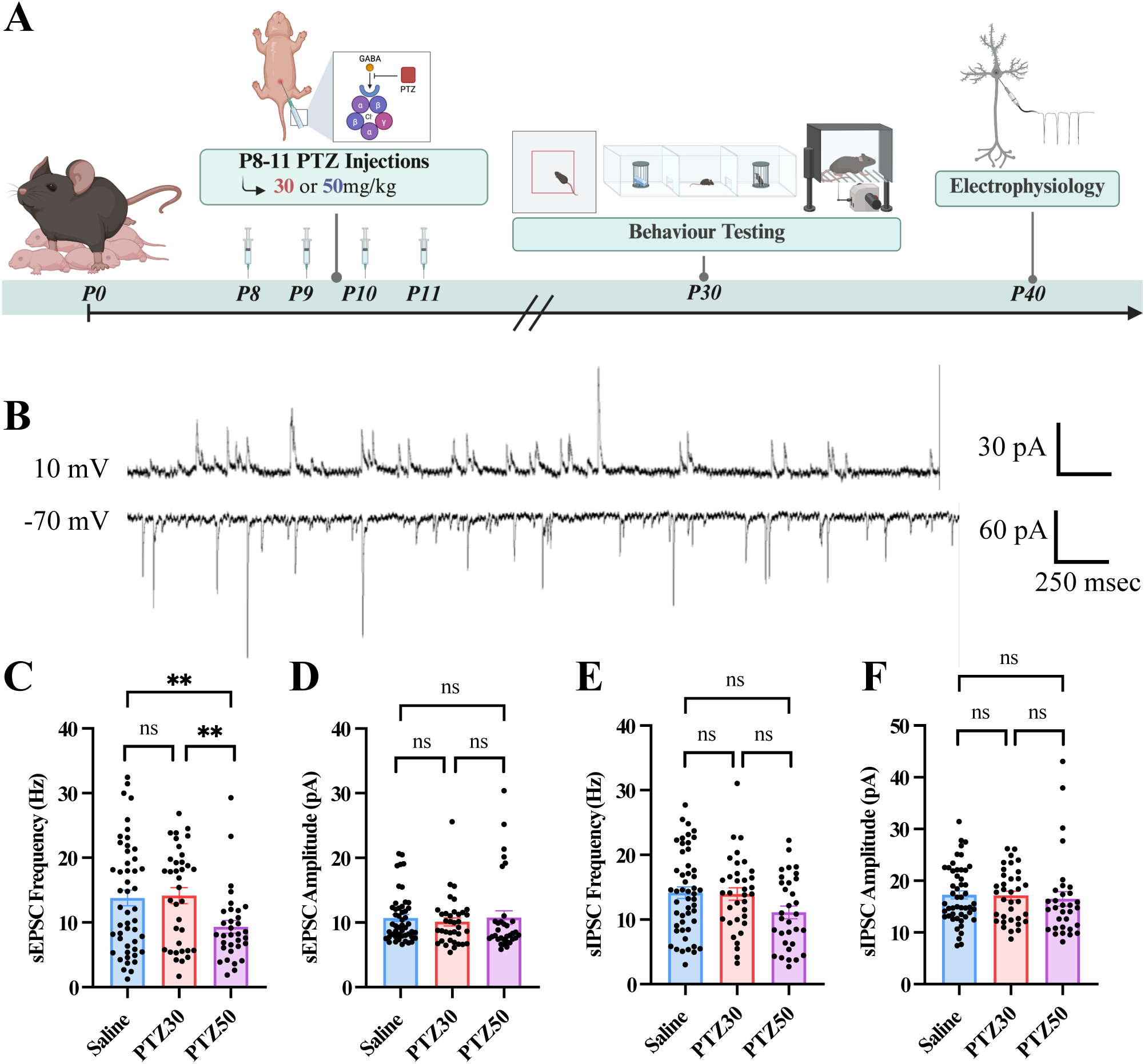
Convulsive PTZ treatment decreases spontaneous excitatory synaptic activity in the ACC. ***A***, Experimental overview of P8-11 PTZ injections, behaviour testing at P30, and electrophysiological recordings around P40. ***B***, Representative whole cell voltage clamp recordings of sIPSCs (top) and sEPSCs (bottom) in L2/3 pyramidal neurons of the ACC from P8-11 PTZ-injected mice at P30. ***C***, 50mg/kg PTZ-injected mice exhibit a significant reduction in sEPSC frequency compared to 30mg/kg and control groups. There were no significant changes in sEPSC amplitude (***D***), sIPSC frequency (***E***), or sIPSC amplitude (***F***) between groups. All figures represented by mean ± SEM. with individual cells as points. P-values were determined by a two-way ANOVA with the Bonferroni correction. **p < 0.01.

We next assessed if the convulsive PTZ dose would induce a similar behavioural phenotype. Similar to our previous experiment, saline-injected mice spent a significantly higher percentage of time interacting with the social stimulus (mouse) than the object (p = 0.0107), whereas PTZ injected mice did not display an interaction preference (Two-way repeated measures ANOVA, effect of stimulus F(1,16) = 9.03, p = 0.0084; interaction effect F(1,16) = 2.40, p = 0.1410; Saline n = 9, PTZ n = 9; Fig.6*A,D*). Similar to the non-convulsive dose, this social preference was observed within the first five minutes of the assay and was abolished for the last five minutes (p = 0.0167), while the 50mg/kg PTZ-treated mice did not display a significant preference (Extended Data Fig. 1B,C). Neither experimental nor control mice displayed a significant interaction preference in the second half of phase two and there were no significant differences in social preferences indices between groups for phase two as a whole or in intervals (Fig. 6*D,H*; Extended Data Fig. 1B,C). Additionally, there were no sex differences in social preference indices between groups (Extended Data Fig. 2C). In phase three, mice injected with a convulsive—50mg/kg—dose of PTZ displayed a significant preference for novel over familiar social interactions in phase three (p = 0.0009). A two-way repeated measures ANOVA revealed a significant interaction effect between stimuli and treatment (F(1,16) = 4.85, p = 0.0426) as well as a significant main effect of stimulus on percentage of time interacting (F(1,16) = 15.87, p = 0.0011; Fig. 6*E*). However, unlike social preference, during the first five minutes of phase three, the novelty preference was significant, but less robust (p = 0.0374) than during the full time. While the main effect of stimulus was still significant for the first five minutes F(1,16) = 8.43, p = 0.0104), the interaction effect was not (F(1,16) = 0.64, p = 0.04366; Extended Data Fig. 1D). The novelty preference index was significantly higher in the PTZ-injected mice compared to controls in phase three (Unpaired Student’s t-test t(16) = 2.422, p = 0.0277; Fig. 6*E*), but not when the phase was divided into the first or second halves (Extended Data Fig. 1D,E). This effect may be driven by a sex difference: PTZ-injected females exhibited a significantly increased novel preference index compared to female controls, as indicated by a significant interaction between stimulus and sex on interaction time (F(1,25) = 4.26, p = 0.0494; Female Saline n = 9, Female PTZ n = 5; Extended Data Fig. 2C). Overall, our data show that convulsive and subconvulsive doses of PTZ between P8-11 induces similar sociability phenotype in the three-chamber test.

**Figure 6.**
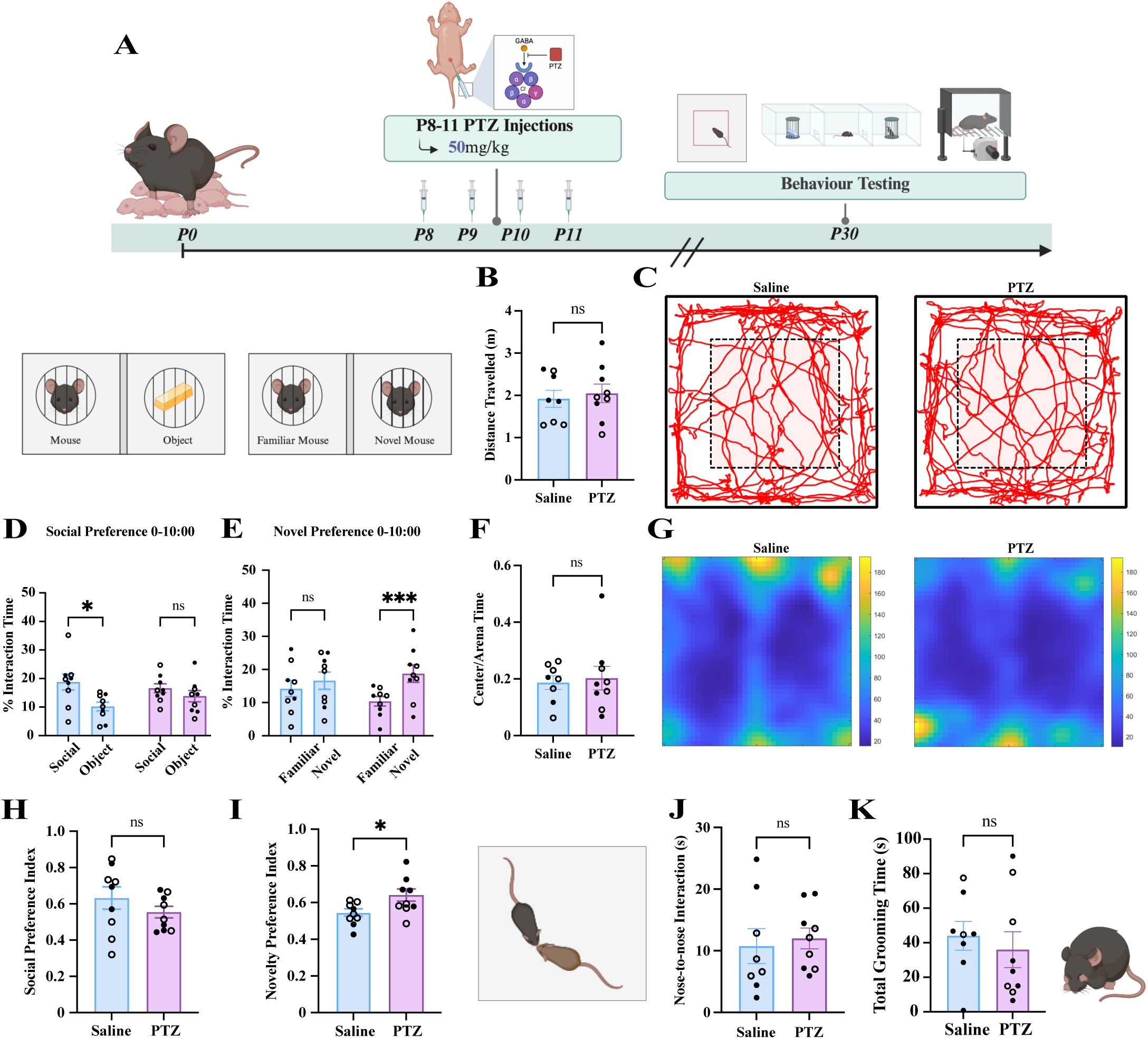
Convulsive PTZ treatment alters social preference without altering anxiety, locomotion, or repetitive behaviour at P30. ***A***, Experimental overview of P8-11 50mg/kg PTZ injections and behaviour testing at P30. ***B***, There were no significant differences in distance travelled between PTZ- or saline-injected mice. ***C,*** Representative tracking of mice the open field arena of saline-(left) and PTZ-injected (right) mice. PTZ-injected mice displayed a lack of significant social preference compared to controls during the full time of phase two (***D***), while the social preference index is unchanged (***H***). PTZ-injected mice displayed a significant novel preference (***E***) and increased novel preference index (***I***) compared to controls during the full time of phase three. ***F***, There were no significant differences in proportion of time spent in the center of the open field arena. ***G,*** Representative heat maps of time spent within the open field arena of saline-(left) and PTZ-injected (right) mice. PTZ-injected mice do not exhibit significant differences in duration of nose-to-nose interactions (***J***) or grooming (***K***) compared to saline-injected mice. Bars represent mean ± SEM. Points represent individual mice. Filled circles represent females while unfilled circles represent males. P-values determined by a two-way ANOVA with the Bonferroni correction or unpaired Student’s t-tests. *p < 0.05, ***p < 0.001.

Following the experimental pipeline of the 30mg/kg PTZ group, the 50mg/kg group was run through the free dyadic social interaction test and the von Frey test (Extended Data Fig. 1G-J). There were no significant differences in withdrawal thresholds between PTZ- or saline-injected mice, however, we observed intra and inter-mouse variability across trials. During the first five minutes of the free dyadic social interaction test–when the test mouse was alone–there were no significant differences between groups in total grooming duration, distance traveled, or proportion of time spent in the center of the arena (Fig. 6 *B*, *F*, *K*; Extended Data Fig. 1H,I). When the stranger mouse was added, we did not observe any significant differences in duration of nose-to-nose interactions or average duration of play behaviour between groups (Fig.6*J*; Extended Data Fig. 1G). There were also no sex differences in any of the previously described behavioural measures. Therefore, like the 30mg/kg group, these data may indicate that convulsive PTZ treatment may selectively impair social preference, as assessed in the three-chamber test.

### Subconvulsive and Convulsive PTZ Treatment from P8-11 Alters Cortical Interneuron Formation

Due to the role of cortical interneurons on regulating excitatory circuits and the role of PVINs in critical period development, we next looked at the potential alterations in PVINs following PTZ injections (Fig. 7*A*). As cortical connectivity and function varies by layer and region, we analyzed interneurons in a layer- and region-specific manner in prelimbic (PL) and infralimbic (IL) areas of mPFC, as well as for the ACC and secondary somatosensory cortical areas (S2) in P40 mice injected with 30mg/kg PTZ. We found layer- and region-specific differences in PVIN and PNN density in the mPFC (two-way repeated measures ANOVA (F(3,21) = 15.10, p < 0.0001; effect of brain region F(3,21) = 23.90, p < 0.0001; Fig. 7*C*). Specifically, PTZ-treated mice exhibited a significant increase in PVINs in L2/3 of the IL region (p = 0.0353) and a significant decrease in PVINs in L5/6 of the PL region (p = 0.0260) compared to controls. We also investigated the presence of perineuronal nets (PNN), a specialized extracellular matrix structure often found surrounding PVINs. In the mPFC, a two-way repeated measures ANOVA showed a significant interaction effect between stimulus and brain region (F(3,21) = 3.32, p = 0.0394), and significant main effects of brain region (F(3,29) = 26.34, p < 0.0001) and treatment, as shown in Figure 7 (F(1,7) = 8.49, p = 0.0226; Fig. 7*C*). Specifically, the PTZ-injected mice displayed a significant increase in PNNs in both L2/3 (p = 0.0040) and L5/6 (p = 0.0158) of the PL regions compared to controls. These findings demonstrate that, like PVINs, PNN development is altered by PTZ-induced hyperexcitation.

**Figure 7.**
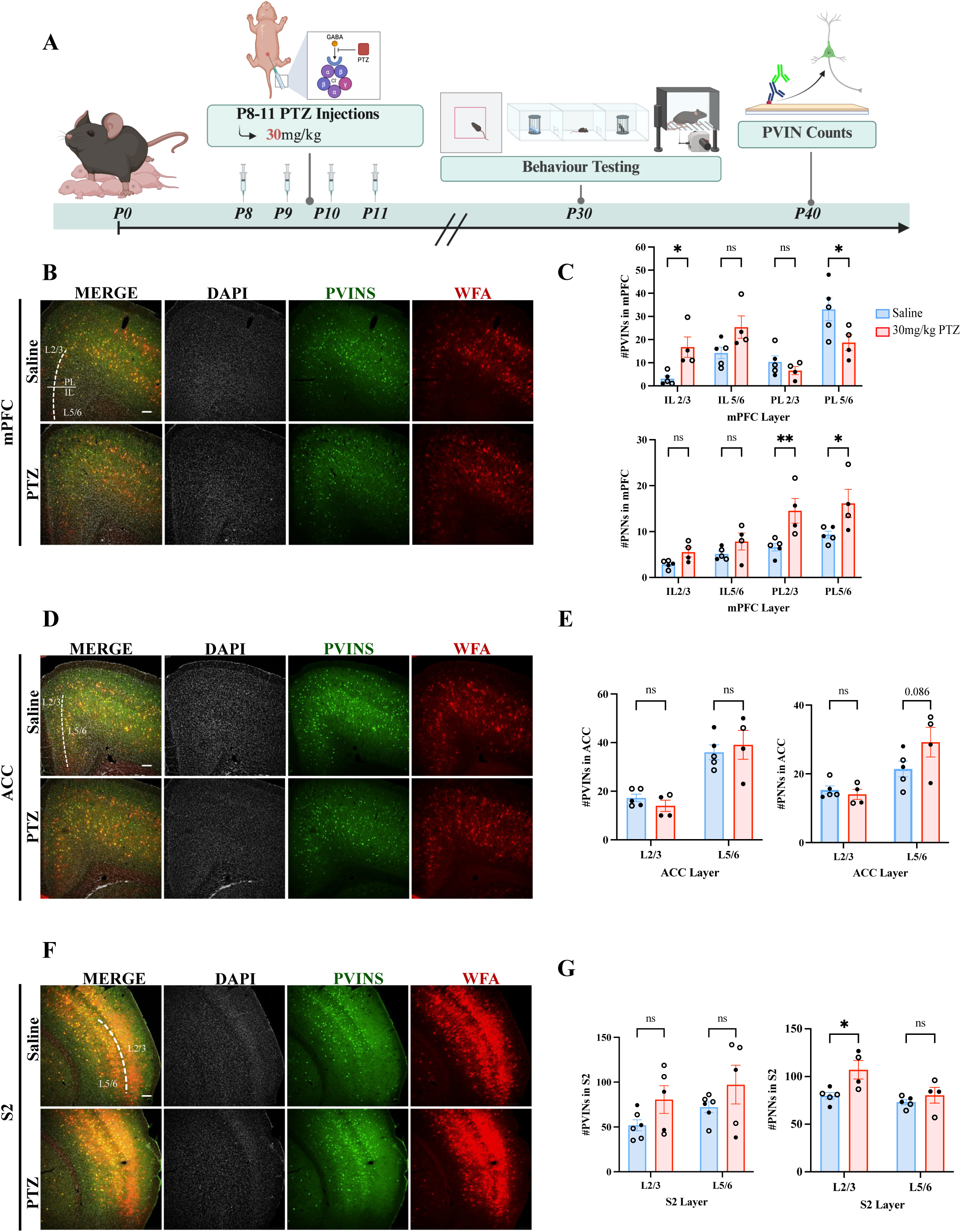
mPFC PVINs alterations following subconvulsive PTZ treatment. ***A***, Experimental overview of PVIN staining following P8-11 PTZ injections and behavioural testing. ***B***, Representative 10x magnification images showing DAPI, PVIN, and WFA staining in the mPFC of saline- and 30mg/kg PTZ-injected mice at P30. ***C***, 30mg/kg PTZ-injected mice had significantly increased PVINs in IL L2/3 and significantly decreased PVINs PL L5/6 (top), as well as significantly increased PNNs in PL L2/3 and L5/6 (bottom). ***D***, Representative 10x magnification images showing DAPI, PVIN, and WFA staining in the ACC of saline- and 30mg/kg PTZ-injected mice at P30. ***E***, There were no significant differences in PVINs (left) or PNNs (right) in the PTZ-injected mice compared to controls. ***F***, Representative 10x magnification images showing DAPI, PVIN, and WFA staining in S2 of saline- and 30mg/kg PTZ-injected mice at P30. ***G***, There were no significant differences in PVINs (left), however, PTZ-injected mice displayed significantly increased PNNs (right) in L2/3. IL = infralimbic, PL = prelimbic. Scale bar is 100 µm. Bars represent mean ± SEM. Points represent average value from three brain slices of a mouse. Filled circles represent females while unfilled circles represent males. P-values were determined by a two-way ANOVA with the Bonferroni correction or unpaired Student’s t-tests. *p < 0.05, **p < 0.01.

We also quantified PVINs and PNNs in ACC (Fig. 7*D*-*E*) and did not observe any significant differences in PVIN quantity between treatment and controls. However, a two-way repeated measures ANOVA revealed a significant main effect of ACC layer on PNN quantity (F(1,7) = 19.4, p = 0.0031; Interaction effect F(1,7) = 3.54, p = 0.102; Fig. *7E*). While there was a trend in increased PNN density in L5/6 of PTZ-treated mice, this was not statistically significant (p = 0.0860). In S2, we observed a significant effect of layer (F(1,7) = 20.9, p = 0.0025) and a significant interaction of layer and treatment (F(1,7) = 7.55, p = 0.0286; Fig7*G*) on PNN density. Specifically, there was a significant increase (p = 0.0144) in the number of PNNs in L2/3 of the 30mg/kg PTZ-injected mice. These results point towards altered development of PNNs in S2 that may warrant further investigation. Finally, we investigated PVINs and PNNs in CPu and CA1 as entire regions (Extended Data Fig. 3). For both regions, there were no significant differences in PVINs or PNNs in the PTZ dosage group compared to controls. Overall, while PNN development was altered by PTZ-treatment in the S2, which has implications with PVIN maturation and stability, our most striking finding was within the mPFC where layer-specific alterations of PVINs and PNNs were observed, suggesting that PTZ-induced hyperexcitation between P8-11 leads to long-term interneuron alterations that are specific to cortical regions directly involved in modulating social behaviour.

To determine if these PTZ-induced alterations in interneurons persisted to adulthood, we quantified PVINs and PNNs following the administration of 30mg/kg PTZ between P8-11. We analysed interneurons in the same regions and using the same methodology in P80-100 mice as previously described. Interestingly, we found significant differences in the number of PNNs, but not PVINs, between PTZ and saline-injected mice at P80 (Extended Data Fig. 4-5). Compared to controls, PTZ-injected mice displayed a significant decrease in the number of PNNs in IL 2/3 (p = 0.0078) but a significant increase in the number of PNNs in PL5/6 (p = 0.0492) (Two-way repeated measures ANOVA, interaction effect F(3,36) = 7.84, p = 0.0004; Brain Region F(3,36) = 5.48, p = 0.0033; n Saline =7, n PTZ = 8; Extended Data Fig. 4C). Interestingly, in the ACC there was a significant decrease (p = 0.0336) in the number of PVINs in L5/6 in the PTZ group compared to controls (Two-way repeated-measures ANOVA, effect of brain region F(1,14) = 5.94: p = 0.0344; effect of treatment F(1,14) = 88.24, p < 0.0001; interaction effect F(1,14) = 0.98; p = 0.3381; n = 14; Extended Data Fig. 4E). Similar quantifications in the S2, CPu and CA1 revealed no significant differences in PVINs or PNNs between groups (Extended Data Fig. 4-5). Taken together, these data indicate that some PTZ-induced changes observed at P40 may be attenuated or shifted by adulthood (P80-100). These temporal differences may shed light on an altered developmental trajectory of PVINs and PNNs following early-life hyperexcitation.

## Discussion

In this study, we demonstrate how increasing neuronal excitability during a sensitive window result in a lasting social behaviour phenotype relevant to ASD. In addition to social preference deficits observed, injection of the GABA_A_ receptor antagonist PTZ of P8-11 in neonatal mice resulted in electrophysiological and histological consequences weeks after the insult. Interestingly, locomotion, anxiety-like behaviour, and mechanical sensitivity were not affected, thus suggesting this critical window is particularly relevant for the development of social behaviour in mice. By employing both non-convulsive and convulsive PTZ doses, we complement previous reports using early-life seizure (ELS) models as well as studies employing chemogenetics or optogenetics to induce non-convulsive developmental hyperexcitation (Bitzenhofer et al., 2021; Castelhano et al., 2013; Medendorp et al., 2021; Lugo et al., 2014). While the degree of hyperexcitation did not appear to produce an alternate sociability phenotype, only convulsive PTZ treatment altered spontaneous cortical network activity. Furthermore, we observed layer- and region-specific alterations in mPFC PVIN number in the 30mg/kg group, which have significant implications for future studies of prosocial circuit function.

### Disrupted Development of Social Preference Following Subconvulsive and Convulsive PTZ

Our work clarifies our understanding of the co-morbidity of ASD and epilepsy in two important ways: 1) seizures are not necessary for developing social preference deficit, and 2) higher cortical excitability during a narrow developmental window is sufficient to disrupt normal development of social preference. Previous studies using early-life hyperexcitation have shown impaired sociability, but whether different excitation levels produce distinct behavioral outcomes remained unclear (Bitzenhofer et al., 2021; Castelhano et al., 2013; Medendorp et al., 2021; Lugo et al., 2014). The administration of PTZ at doses shown to be non-convulsive (30mg/kg) and convulsive (50mg/kg) in previous research resulted in a similar phenotype, indicating that convulsions are not required (Monteiro et al., 2024; Parker et al., 2016; Shimada and Yamagata 2018). Indeed, we found that PTZ-injected mice of the 30mg/kg and 50mg/kg groups did not exhibit a significant interaction preference throughout phase two of the three-chamber test at P30, a time point when wildtype mice displayed social preference, which is absent in the *Shank3b*^-/-^ ASD model (Balaan et al., 2019; Fairless et al., 2019). This social preference in saline-injected mice was strongest during the initial minutes, which is consistent with previous reports (Fairless et al., 2019; Moy et al., 2004; Nadler et al., 2004; Page et al., 2009; Selimbeyoglu et al., 2017). Although the sociability deficits were present in both PTZ doses, the concentrations used are relatively mild compared to doses used in epilepsy models, upwards of 80-100mg/kg (Monteiro et al., 2024; Parker et al., 2016). Therefore, the fact that only four daily injections of 30mg/kg PTZ results in a reduced social preference index and nose-to-nose interactions is quite remarkable. The observed phenotype in our PTZ model aligns with findings of previous research showing that *Scn1a^+/−^,* BTBR, and C58J mouse models of ASD display fewer interactions in the reciprocal interaction test (Chang et al., 2018; Han et al., 2012; Silverman et al., 2015; Yang et al., 2012), raising the possibility that sociability phenotype in these mice is determined by altered neuronal development during this short P8-11 sensitive window.

Given the effects of PTZ on social preference and nose-to-nose interactions, we were surprised by the absence of effects of PTZ on the duration of juvenile play which is often used as an additional index of reciprocal social interactions in developing mice (Silverman et al., 2010; Terranova and Laviola, 2005). However, juvenile play is inconsistently impacted across mouse models of ASD and thus might not mirror social preference as measured in the three-chamber test (Bey and Jiang 2015; McFarlane et al., 2007; Penagarikano et al., 2011). Since there were no significant differences in the duration of play behaviour between any treatment or control mice, the effects of PTZ may be specific to the development of social preference, and potentially direct social interactions rather than other behaviours related to particular aspects of sociability or interindividual communication. However, it is worth noticing that the recommended duration of the juvenile play test is 10 minutes (Terranova and Laviola, 2005); therefore, our shorter window of five minutes may not have been sufficient to detect alterations in play behaviour.

Unlike social preference, social novelty preference as measured in phase 3 of the three-chamber test, was largely absent across most groups including saline-injected controls. This result was unexpected, as adult mice typically prefer novel over familiar conspecifics, a preference often lacking in ASD models (Chang et al., 2018; Ey et al., 2013; Han et al., 2012; Yang et al., 2012). One explanation is that P30 might be too early for this preference to occur, as social novelty preference has only been reported in mice that were at least six weeks old (Moy et al., 2004). It is also worth mentioning that social novelty measure could also be impacted by a putative deficit in social memory (Leung et al., 2018). Indeed, social novel preference requires proper memory retrieval to discriminate between the conspecific from phase two and the novel conspecific (Gunaydin et al., 2014; Leung et al., 2018). Therefore, determining if P30 mice have the ability to form and retrieve social memory would be needed to understand the absence of social novelty preference observed.

### P8-11 PTZ Treatment Does not Disrupt Grooming, Anxiety, Locomotion, or Mechanical Sensitivity

An interesting finding from our experiments is that PTZ injection between P8-11 disrupts social preference specifically, without altering anxiety-like, repetitive, or locomotor behaviour. This finding is particularly relevant with regard to self-grooming, a behaviour sometimes affected in autism models which is thought to reflect the increased repetitive behaviours observed in ASD (Silverman et al., 2010). However, we did not observe differences in self-grooming behaviour between PTZ-injected and control mice (Kazdoba et al., 2016; Silverman et al., 2010). Interestingly, these findings align with Bitzenhofer and coauthors (2021) who found no differences in self-grooming behaviour between mice who underwent optogenetic stimulation of the mPFC from P7-11. This, along with the lack of hypo- or hyperactivity, may be due to P8-11 not overlapping with the maturation of striatal circuitry involved in repetitive behaviours and goal-oriented locomotion behaviours (Kazdoba et al., 2016; Peixoto et al., 2016, 2019). This raises the possibility that, at least in mice, behavioural alterations observed in ASD models do not arise from perturbation during the same sensitive developmental window.

We investigated anxiety-like behaviour since anxiety is known to be more common in youth with ASD compared to their peers (Avni et al., 2018; Wood et al., 2010). As a measure of anxiety-like behaviour, we used the time spent in the center of an open field arena (Seibenhener and Wooten, 2015). There were no significant differences in anxiety between the PTZ- and saline-injected mice from either dosage group, which aligns with the findings of Bitzenhofer et al., (2021). It is, however, important to note that the comorbidity between anxiety and ASD is complex and—like other ASD-relevant behaviours—cannot be fully untangled in rodent studies (Ey et al., 2011). In the context of social behaviour, symptoms of anxiety may overlap with symptoms ASD symptoms such as anxiety surrounding social fearfulness increasing social withdrawal behaviour (Kerns et al., 2014).

Finally, the von Frey test was used to assess mechanical sensitivity as unusual somatosensory responsiveness—including hypo- and hypersensitivity—has been observed in individuals and animal models of ASD (Bogdanova et al., 2022; Deuis et al., 2017). Analysis of paw withdrawal thresholds revealed no significant differences between PTZ groups of either dosage. As mechanical sensitivity was not addressed in previous developmental hyperexcitation studies, we provide preliminary evidence that neither subconvulsive nor convulsive hyperexcitation from P8-11 impacts the development of mechanical sensitivity (Bitzenhofer et al., 2021; Castelhano et al., 2013; Lugo et al., 2014; Medendorp et al., 2021). In sum, our data strongly suggest that the sociability deficit observed following PTZ is neither caused by gross motor or somatosensory impairments, nor confounded by anxiety-like phenotype.

### Lasting Effects of Transient PTZ Treatment on Cortical Excitability and Interneuron Maturation

An interesting feature of our model is that four daily injections of a low dose of PTZ have long-lasting consequences on normal brain development that parallels the observed behavioural phenotype. For instance, P8-11 50 mg/kg PTZ injections resulted in a significant reduction in the frequency of spontaneous excitatory postsynaptic currents (sEPSCs), suggesting decreased spontaneous excitatory cortical network activity. This decrease in sEPSC frequency is interesting in the context of the increased in cortical cFos+ nuclei at P11, likely reflecting compensatory mechanisms regulating cortical network activity in an effort to maintain E/I balance via homeostatic plasticity (Turrigiano et al., 2012). This can occur through a variety of physiological mechanisms and impaired homeostatic plasticity responses have been characterized in mouse models of ASD (Antoine et al., 2019; Del Pino et al., 2018; Nelson and Valakh, 2015). Given there were no significant changes in spontaneous network activity in the 30mg/kg group at P30, this network adaptation may not have occurred in the subconvulsive group and thus might not be underlying the behavioural phenotype.

We also investigated PVINs which are critical to maintain proper E/I ratio and are affected in various ASD models (Filice et al., 2020). An important limitation in our PVIN quantifications is that we did not investigate potential differences in the expression of PV itself. From our data, layer- and region-specific alterations in mPFC PVIN may be pertinent to the observed social behaviour phenotype given the importance of the mPFC in mouse social behaviour (Sato et al., 2023). For example, the increased number of PVINs in the IL following the 30mg/kg P8-11 PTZ treatment could translate to reduced activity of the prosocial pathway innervating the nucleus accumbens (Park et al., 2021). However, future experiments targeting or rescuing specific circuits would be required to determine the neuronal pathway affected by PTZ injection.

The PVINs changes are interesting in the context of our electrophysiological data as there were no significant differences in sEPSCs or sIPSCs in pyramidal neurons in the ACC in the 30mg/kg group compared to controls. Therefore, assessing the electrophysiological properties of PVINs in the mPFC is warranted to determine if an increase in PVINs translates to more inhibitory firing and may be relevant to our observed social behaviour phenotype. This is particularly important as Bitzenhofer et al., (2021) found increased PVINs in the mPFC, but determined these were hypofunctional at P37-40. Despite the PVIN alterations at P30, we did not observe differences in PVIN numbers between the 30mg/kg group and controls at P80. It is possible that the changes attenuated or resulted from an adaptation of the interneuron network in the mPFC (Contractor et al., 2021; Donato et al., 2013). Despite being past the window of PVIN apoptosis, it is also possible that the P30 PVIN changes were transient, given P30 is a time window important for synaptic pruning and refinement of circuitry (Semple et al., 2013; Lim et al., 2018; Wong et al., 2018).

The increased PNNs in the PL regions of the 30mg/kg group warrants discussion given PNNs are crucial for PVIN stability and critical window timing (Cisneros-Granco et al., 2020). For example, if aberrantly formed, PNNs could induce a premature loss of plasticity or closure of critical windows. Interestingly, at P80, 30mg/kg PTZ-injected mice exhibited an alternate pattern of PNN distribution with decreased PNNs in the IL and increased PNNs in PL. Since these regions appear to have opposite in sociability (Sato et al., 2023), it raises the possibility that neuronal activity and interneuron maturation in these regions be inversely correlated. Although the impact of these PNNs changes is unclear, it is known that PNNs are still developing at P30 and it is therefore not surprising that the situation is different at P80 (Lupori et al., 2023; Mirzadeh et al., 2019).

PVINs in the ACC were investigated as they modulate processes such as attention and empathy relevant to sociability (Kietzman et al., 2023). While models of ASD exhibit reduced PVINs in the ACC there were no significant differences in PVINs between the PTZ dosage groups at P30 (Filice et al., 2020). Therefore, PVIN changes in our PTZ model at P30 may be localized to the more rostral IL and PL mPFC, which is further corroborated by the lack of PVIN differences observed in S2. Finally, we assayed PVINs in non-cortical regions including CA1 and the CPu, which are relevant to previous studies using in ELS and *Shank3b*^-/-^ models, respectively (Bernard et al., 2015; Castelhano et al., 2013; Peixoto et al., 2016, 2019). We did not find any significant differences in PVINs or PNNs in CA1 or the Cpu in either PTZ treatment group at P30 or P80. In summary, our PVIN data suggest that PTZ treatment from P8-11 induces dose-dependent alterations to PVINs in mPFC regions relevant to social behaviour that are dosage specific. Interestingly, these changes may persist to adulthood and future studies will be needed to determine how the observed PVINs changes lead to functional alterations in the mPFC.

Overall, we found that P8-11 represents a critical window in which the formation of social preference was susceptible to hyperexcitation. Mice injected with non-convulsive or convulsive doses of PTZ from P8-11 exhibited a lack of social preference at P30. In contrast, the effects of PTZ dosage on cortical activity and PVIN formation may suggest that distinct mechanisms are implicated in the shared behavioural phenotype. Ultimately, this model opens the door to investigate alterations in sociability-related pathways that may mediate behavioural and neuronal phenotypes relevant to both ASD and epilepsy.

---

**Extended Figure 1.**
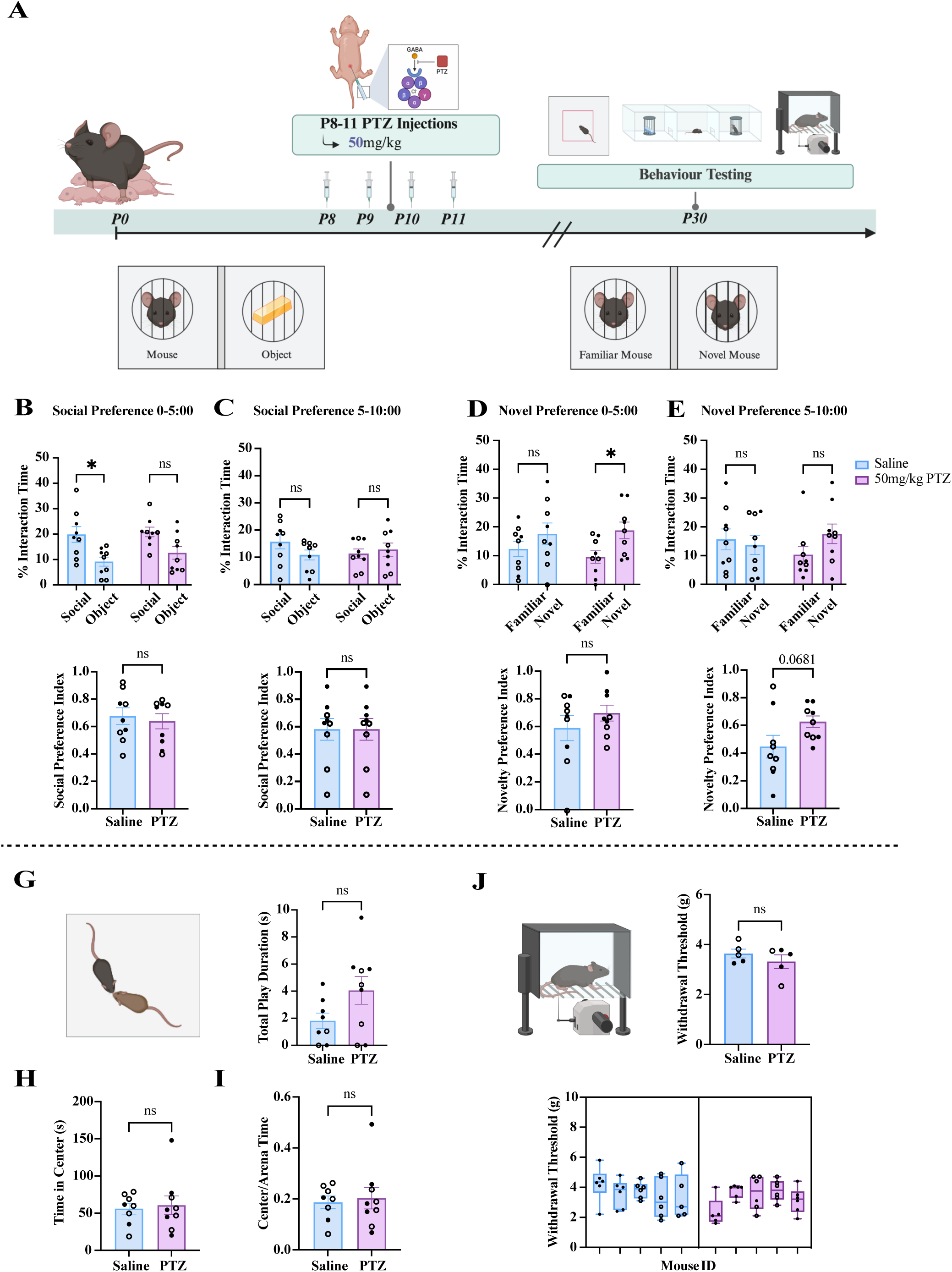
50mg/kg P8-11 PTZ treatment alters social preference in the three-chamber test at P30. ***A***, Experimental overview of P8-11 30mg/kg PTZ injections and behaviour testing at P30. PTZ-injected mice displayed a lack of significant social preference compared to controls during the first half (***B***), but not last half (***C***) of phase two. PTZ-injected mice displayed a significant novel preference compared to controls during the first half (***D***), but not last half (***E***) of phase three. ***F***, PTZ-injected mice do not display significant difference in grooming time of time compared to controls. ***G***, There were no significant differences in duration of play behaviour between treatment and controls. PTZ-injected mice do not display significant difference in total time (***H***) or proportion of time (***I***) spent in the center of the open field arena. ***J***, There were no significant differences in withdrawal thresholds on the Von Frey test between treatments and controls. Bars represent mean ± SEM. Points represent individual mice. Filled circles represent females while unfilled circles represent males. P-values determined by a two-way ANOVA with the Bonferroni correction or unpaired Student’s t-tests. *p < 0.05.

**Extended Figure 2.**
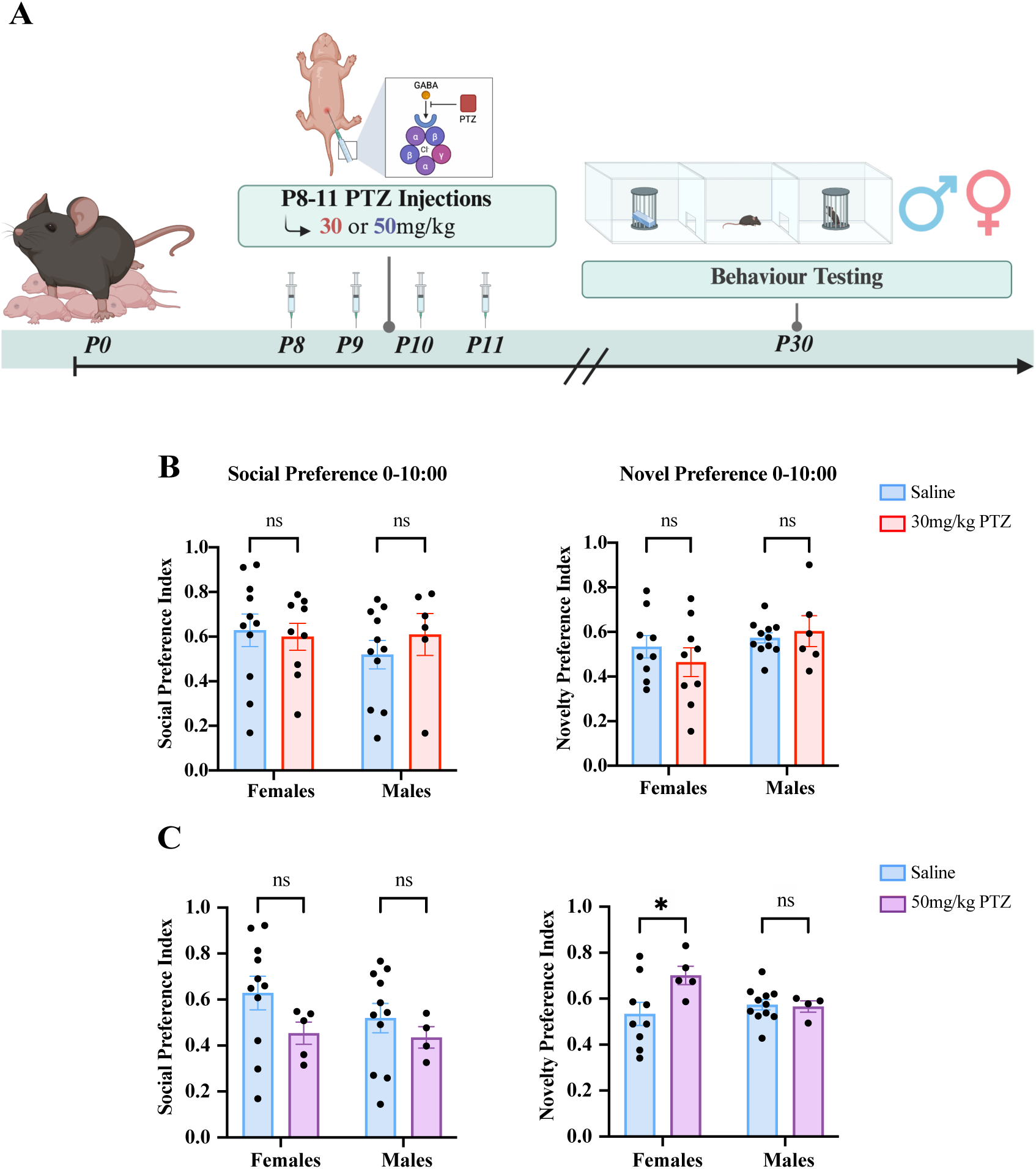
Sex-specific analysis of social and novel preference indices in the three-chamber test for 30 and 50mg/kg P8-11 PTZ-injected mice at P30. ***A***, Experimental overview of P8-11 30 or 50mg/kg PTZ injections and behaviour testing at P30. ***B***, Neither the 30mg/kg PTZ- or saline-injected mice do not display sex-specific differences in social (left) or novel (right) preference in the three-chamber test. ***C***, 50mg/kg PTZ-injected mice do not display sex differences in social preference (left), but female 50mg/kg PTZ-injected mice display a significantly increased novel preference compared to female controls. Points represent individual mice. P-values determined by a two-way ANOVA with the Bonferroni correction. *p < 0.05.

**Extended Figure 3.**
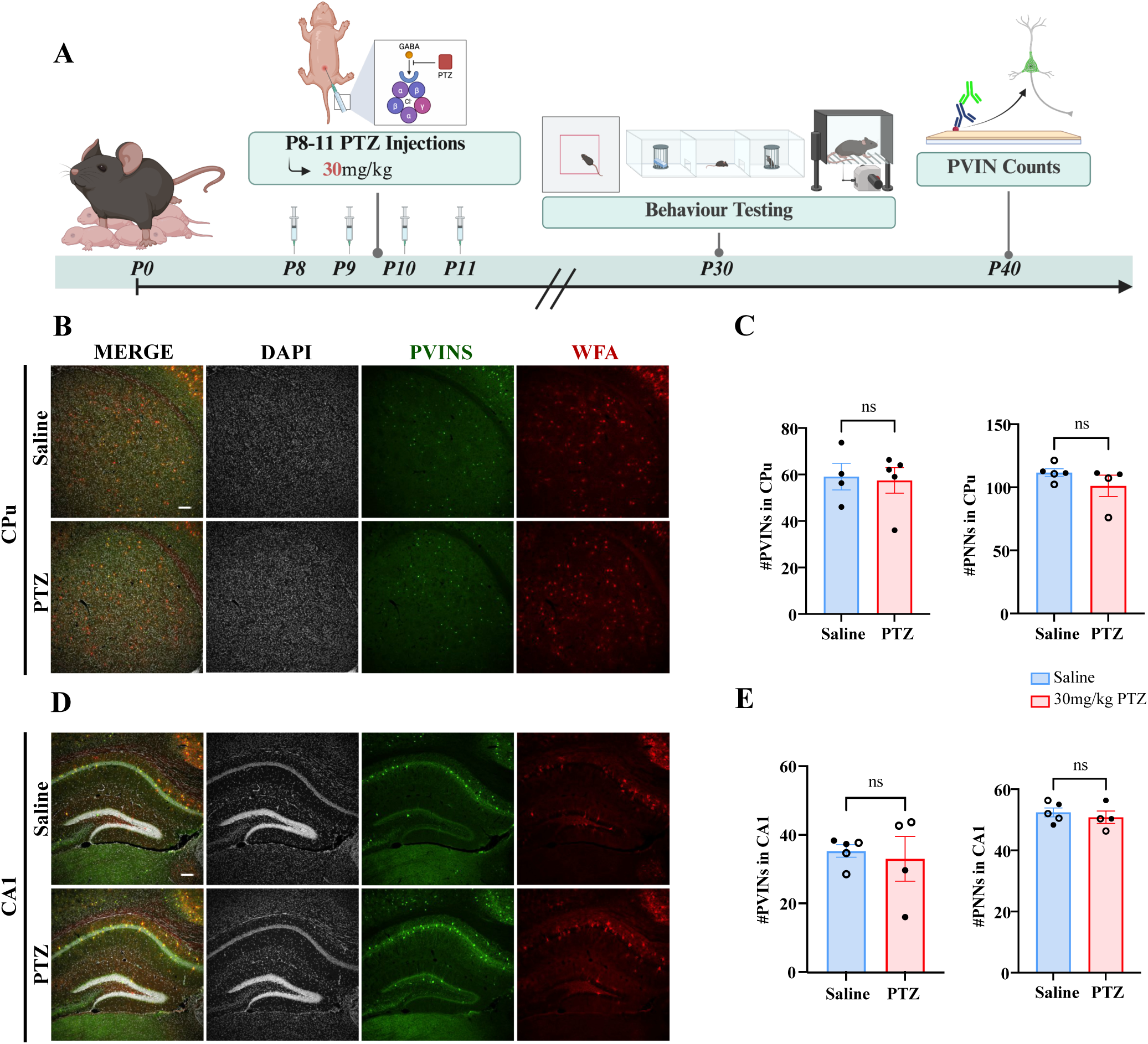
Non-cortical PVINs are unaltered in 30mg/kg PTZ-injected mice at P30. ***A***, Experimental overview of PVIN staining following P8-11 PTZ injections and behavioural testing. ***B***, Representative 10x magnification images showing DAPI, PVIN, and WFA staining in the CPu of saline- and 30mg/kg PTZ-injected mice at P30. ***C***, There were no significant differences in PVINs (left) in or PNNs (right) of PTZ-injected mice compared to controls. ***D***, Representative 10x magnification images showing DAPI, PVIN, and WFA staining in CA1 of saline- and 30mg/kg PTZ-injected mice at P30. ***E***, There were no significant differences in PVINs (left) in or PNNs (right) of PTZ-injected mice compared to controls. Scale bar is 100 µm. Bars represent mean ± SEM. Points represent average value from three brain slices of a mouse. Filled circles represent females while unfilled circles represent males. P-values were determined by unpaired Student’s t-tests.

**Extended Figure 4.**
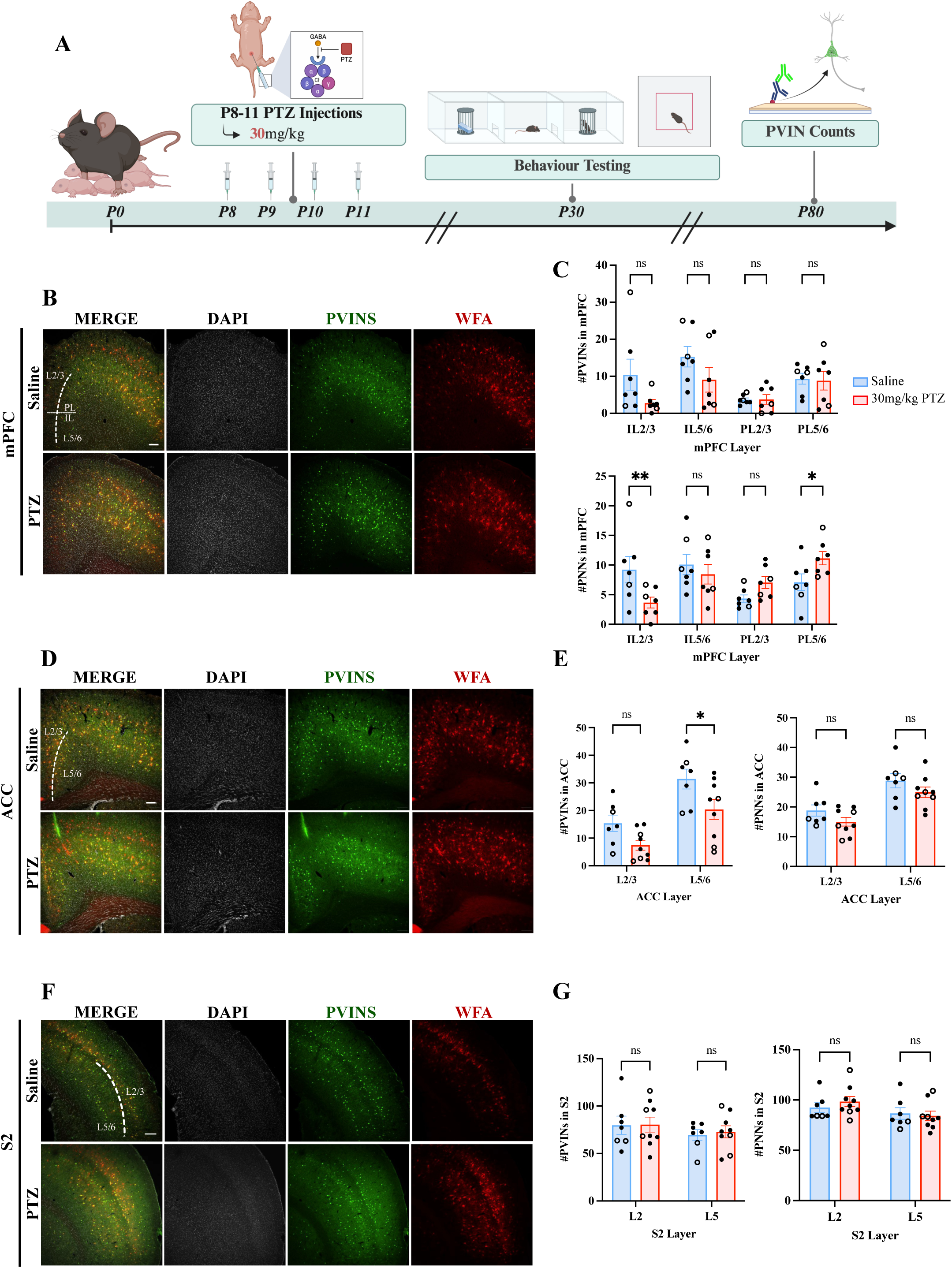
mPFC PNNs are differentially altered in 30mg/kg PTZ-injected mice at P80. ***A***, Experimental overview of PVIN staining following P8-11 PTZ injections and behavioural testing. ***B***, Representative 10x magnification images showing DAPI, PVIN, and WFA staining in the mPFC of saline- and 30mg/kg PTZ-injected mice at P80. ***C***, 30mg/kg PTZ-injected mice did not display significant differences in PVINs compared to controls (top), however, PTZ-injected mice had significantly decreased PNNs in IL L2/3 and significantly increased PNNs PL L5/6 (bottom). ***D***, Representative 10x magnification images showing DAPI, PVIN, and WFA staining in the ACC of saline- and 30mg/kg PTZ-injected mice at P80. ***E***, There were no significant differences in PVINs (top) or PNNs (bottom) in the PTZ-injected mice compared to controls. ***F***, Representative 10x magnification images showing DAPI, PVIN, and WFA staining in S2 of saline- and 30mg/kg PTZ-injected mice at P30. ***G***, There were no significant differences in PVINs (left), however, PTZ-injected mice displayed significantly increased PNNs (right) in L2/3. IL = infralimbic, PL = prelimbic. Scale bar is 100 µm. Bars represent mean ± SEM. Points represent average value from three brain slices of a mouse. Filled circles represent females while unfilled circles represent males. P-values were determined by a two-way ANOVA with the Bonferroni correction or unpaired Student’s t-tests. *p < 0.05, **p < 0.01.

**Extended Figure 5.**
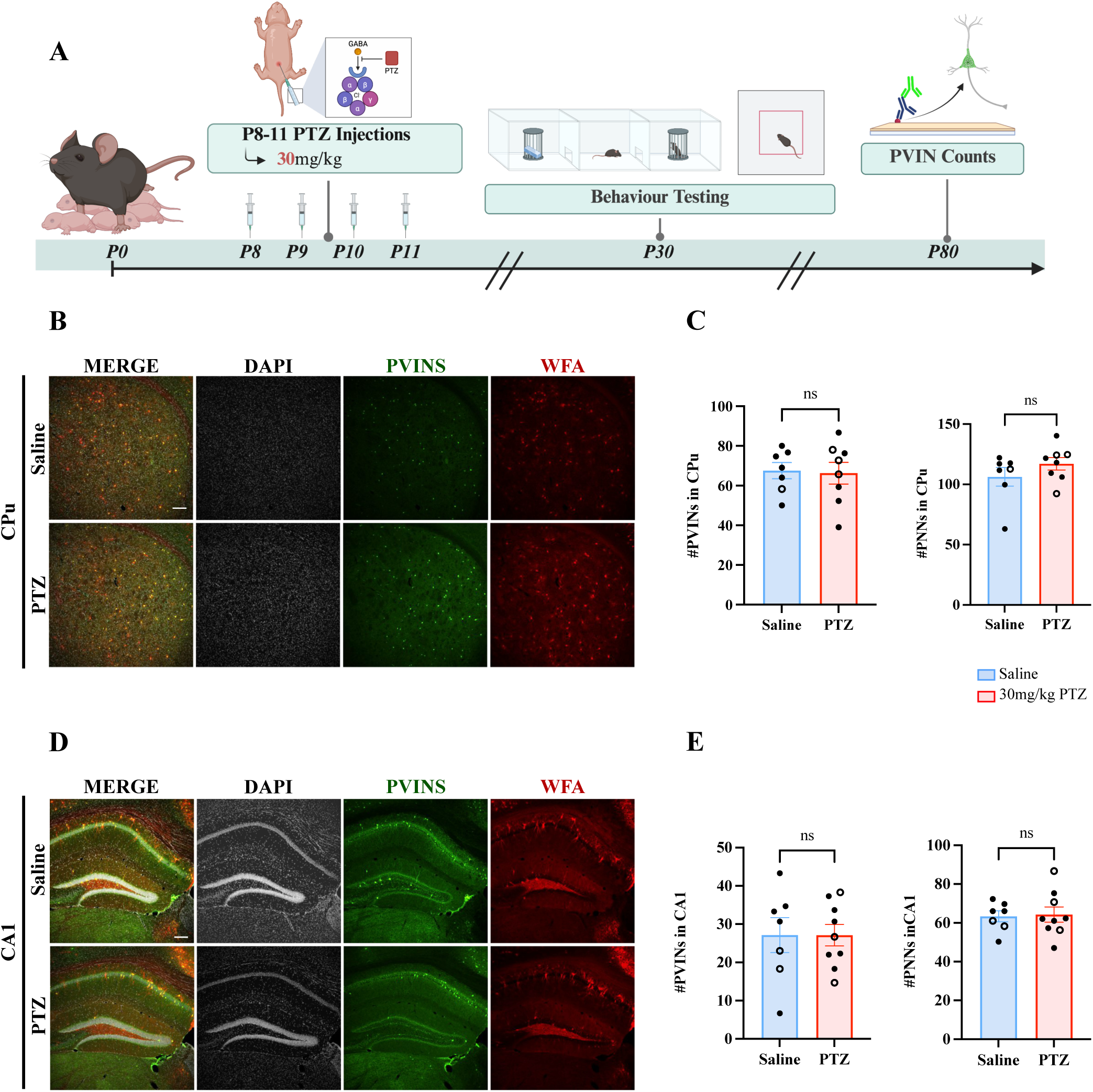
Non-cortical PVINs are unaltered in 30mg/kg PTZ-injected mice at P80. ***A***, Experimental overview of PVIN staining following P8-11 PTZ injections and behavioural testing. ***B***, Representative 10x magnification images showing DAPI, PVIN, and WFA staining in the CPu of saline- and 30mg/kg PTZ-injected mice at P80. ***C***, There were no significant differences in PVINs (top) in or PNNs (bottom) of PTZ-injected mice compared to controls. ***D***, Representative 10x magnification images showing DAPI, PVIN, and WFA staining in CA1 of saline- and 30mg/kg PTZ-injected mice at P80. ***E***, There were no significant differences in PVINs (top) in or PNNs (bottom) of PTZ-injected mice compared to controls. Scale bar is 100 µm. Bars represent mean ± SEM. Points represent average value from three brain slices of a mouse. Filled circles represent females while unfilled circles represent males. P-values were determined by unpaired Student’s t-tests.

## Notes

### Competing Interest Statement

The authors have declared no competing interest.

### Funding

- Azrieli Foundation

